# A review of models of natural pest control: toward predictions across agricultural landscapes

**DOI:** 10.1101/2020.03.13.990531

**Authors:** Nikolaos Alexandridis, Glenn Marion, Rebecca Chaplin-Kramer, Matteo Dainese, Johan Ekroos, Heather Grab, Mattias Jonsson, Daniel S. Karp, Carsten Meyer, Megan E. O’Rourke, Mikael Pontarp, Katja Poveda, Ralf Seppelt, Henrik G. Smith, Emily A. Martin, Yann Clough

## Abstract

Natural control of crop pests has the potential to complement or replace intensive agricultural practices, but its mainstream application requires reliable predictions in diverse socioecological settings. In lack of a widely accepted model of natural pest control, we review existing modelling approaches and critically examine their potential to provide understanding and predictions across agricultural landscapes. Models that explicitly represent the underlying mechanisms are better positioned to represent the diversity and context sensitivity of natural pest control than correlative models. Such mechanistic models have used diverse techniques to represent crop-pest-enemy combinations at various spatiotemporal scales. However, certain regions of the world and socioeconomic aspects of natural pest control are underrepresented, while modelling approaches are restricted by a fundamental trade-off between generality and realism. We propose that modelling natural pest control across agroecosystems requires a framework of context-specific generalizations, based on empirical evidence and theoretical expectations. Reviewed models of natural pest control indicate potential attributes of such a general predictive framework.

## Introduction

A substantial portion of crops around the world is consumed by invertebrate pests, with resulting yield losses predicted to increase in response to climate warming (Deutsch et al., 2018). Control of crop pests by their natural enemies, such as arthropod predators and parasitoids, is an essential ecosystem service valued over a decade ago at US$4.5 billion/yr in the USA (Losey & Vaughan, 2006). Natural pest control holds the potential to at least partially replace intensive agricultural practices aimed at pest regulation, including wide-spread pesticide use (Khan et al., 2014; Tschumi et al., 2015; Holland et al., 2017). However, mainstreaming natural pest control in agriculture requires simple and reliable tools that can predict its response to land management across agroecosystems (Kleijn et al., 2019).

Applicable tools already exist to map a range of ecosystem services on the basis of detailed land-use maps, including crop pollination by invertebrates (Sharp et al., 2014). Similar to pollinators, the abundance and diversity of natural enemies and their potential to provide ecosystem services are also affected by agricultural land use (Tscharntke et al., 2005). In contrast to pollination, natural pest control involves an additional trophic level, while pest and enemy biology and the associated impacts on crops can be extremely diverse across agroecosystems (Tscharntke et al., 2016; Karp et al., 2018). This has restricted the potential to link specific landscape characteristics to enhanced pest suppression across cases (Englund et al., 2017). As a result, efforts to assess natural pest control on the basis of spatially-explicit land-use information have had limited predictive scope (e.g., Rega et al., 2018). Currently, no widely accepted model for assessment of the natural pest control potential of agricultural landscapes exists. In addition, the usefulness of existing models outside the socioecological settings and purpose for which they were developed is unknown.

Our objective is to evaluate the ability of existing modelling frameworks to predict the responses of natural pest control to land-use changes across agricultural landscapes. We review models of natural control of invertebrate pests, examining the strengths and weaknesses of modelling approaches and techniques. We focus on the systems and processes that are represented by existing models of natural pest control, as well as the main properties of model output. We use the results of the review to indicate promising directions and potential attributes of models predicting natural pest control responses to agricultural management across agroecosystems.

### Literature search

A search of the Core Collection of ISI Web of Science with the topic argument: model* AND land* AND (biocontrol OR “biological control” OR “pest control” OR “natural control”) returned 448 publications (search date: 25 June 2018). We reviewed their abstracts, discarding non-modelling studies as well as studies of non-crop systems, anthropogenic biocontrol, and plant or vertebrate pests. We thus retained 172 studies modelling natural control of invertebrate pests in agricultural systems. Of the respective models, 105 are formulated on the basis of observed correlations among system components (correlative) and 67 are based on the representation of explicit causative agents (mechanistic) (Fig. 1a). Mechanistic approaches dominated the early models of natural pest control; however, the picture has been reversed in recent years (Fig. 1c).

**Fig. 1.**
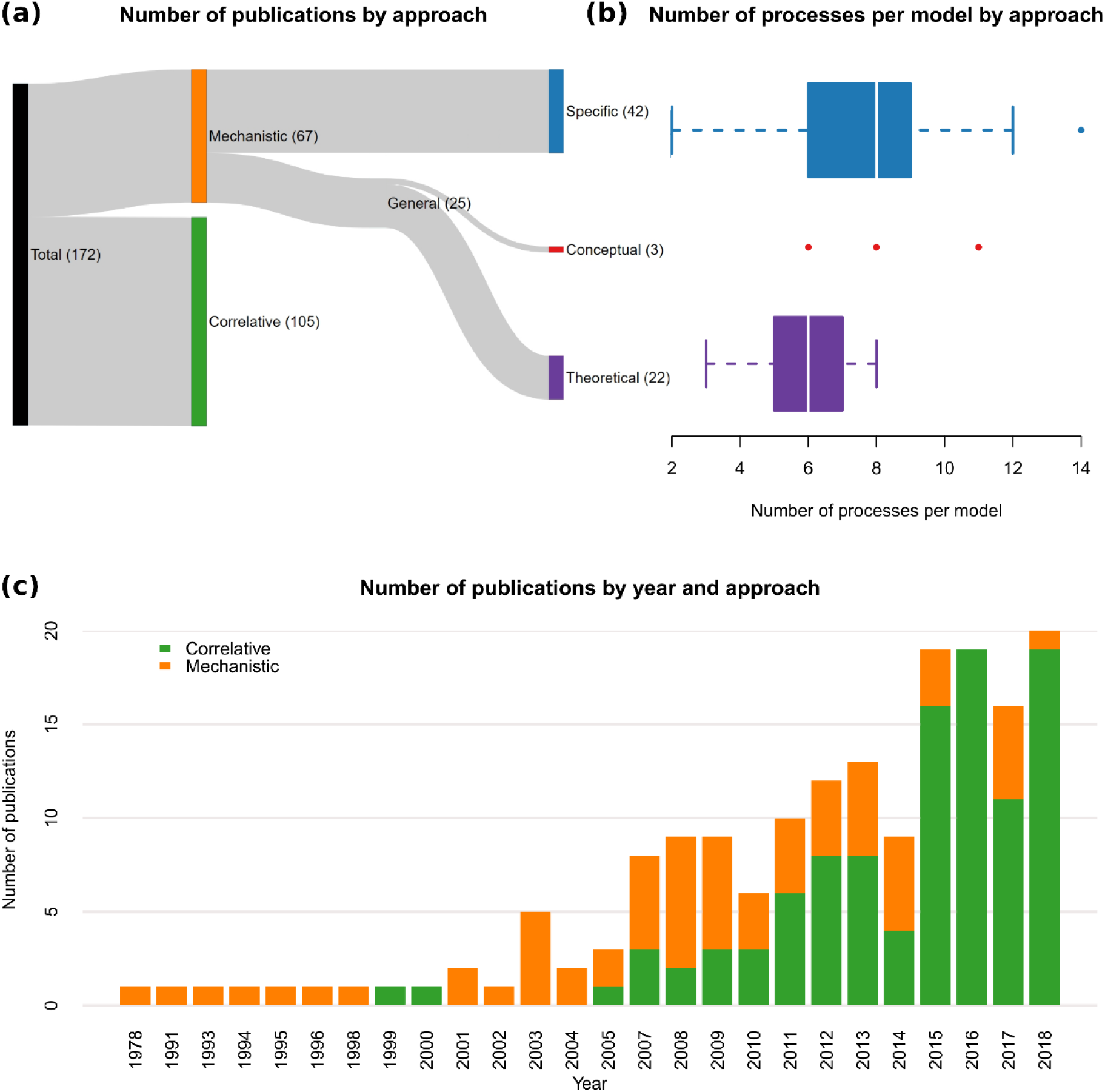
(a) Number of reviewed publications following contrasting modelling approaches, sequentially separated based on their mechanistic basis, generalization potential and approach to generality. (b) Number of processes represented per reviewed model, among models following three contrasting mechanistic approaches. Box bands, lower and upper edges represent 2^nd^, 1^st^ and 3^rd^ quartiles, respectively. Whiskers extend to the most extreme values within 1.5 × interquartile range from the box; more extreme values are indicated with dots. (c) Number of reviewed publications featuring correlative and mechanistic models for each publication year.

### Correlative models

The main objective of correlative models is the assessment and development of general theories explaining the responses of natural pest control to agricultural land use at scales ranging from field to region. The majority of these models are applied at the scale of landscape or higher. A fundamental theoretical expectation is that compositional (e.g., proportion of non-crop habitat) and/or configurational complexity (e.g., field edge density) of the landscape surrounding a field enhances in-field activity of natural enemies by providing them with limiting resources, such as nesting sites and alternative food sources (Landis et al., 2000). At the same time, landscape simplification is generally expected to enhance agricultural pests by increasing the density of their host plants (Bianchi et al., 2006).

Meta-analyses of empirical studies show positive effects of landscape complexity on the diversity and abundance of natural enemies but, critically, the responses of pest suppression and crop yield are rather inconsistent (Bianchi et al., 2006; Chaplin-Kramer et al., 2011). Analyses of recent correlative models extending to continental and global scales corroborate the inconsistent nature of natural pest control responses to landscape characteristics. These studies identified systematic (Karp et al., 2018) and trait-mediated differences in responses (Martin et al., 2019) as a major underlying cause of discrepancies.

As a general understanding of natural pest control currently eludes us, the use of correlative models for predictive purposes requires the development of models for every system of interest. This task involves extensive collection of data, including an array of land-use, agronomic, abiotic and biotic environmental variables, hindering the derivation of accurate predictions across agricultural landscapes. Furthermore, correlative models typically make several assumptions, such as stationarity of ecological processes or lack of adaptability, that may not hold in novel or non-equilibrium contexts often associated with environmental change (Dormann, 2007).

### Mechanistic models

Models that explicitly represent causative agents of ecological observations allow for more reliable extrapolations in space and time than purely correlative approaches (Gotelli et al., 2009). Deviations of predictions from observations can generate knowledge about the role of specific processes and indicate the most productive areas for future research (Soetaert & Herman, 2009). However, deep knowledge on system function, which is required for the formulation of these mechanistic models, excludes less-studied systems, while data requirements for model parameterization and validation can prove just as challenging as for correlative approaches (Dormann et al., 2012). Furthermore, although model complexity may be essential for precise and accurate predictions, it can also restrict a model’s transferability (Yates et al., 2018). It is not surprising that the general trend among mechanistic models of natural pest control toward more processes per model has been checked by attempts at simpler system representations (Fig. 1b, 2a). Still, inconsistencies in the responses of natural pest control and our limited understanding of the underlying causes favour the use of mechanistic models for the generation of knowledge and its reliable transfer between agroecosystems. We therefore examine in more detail the characteristics of mechanistic models of natural pest control.

**Fig. 2.**
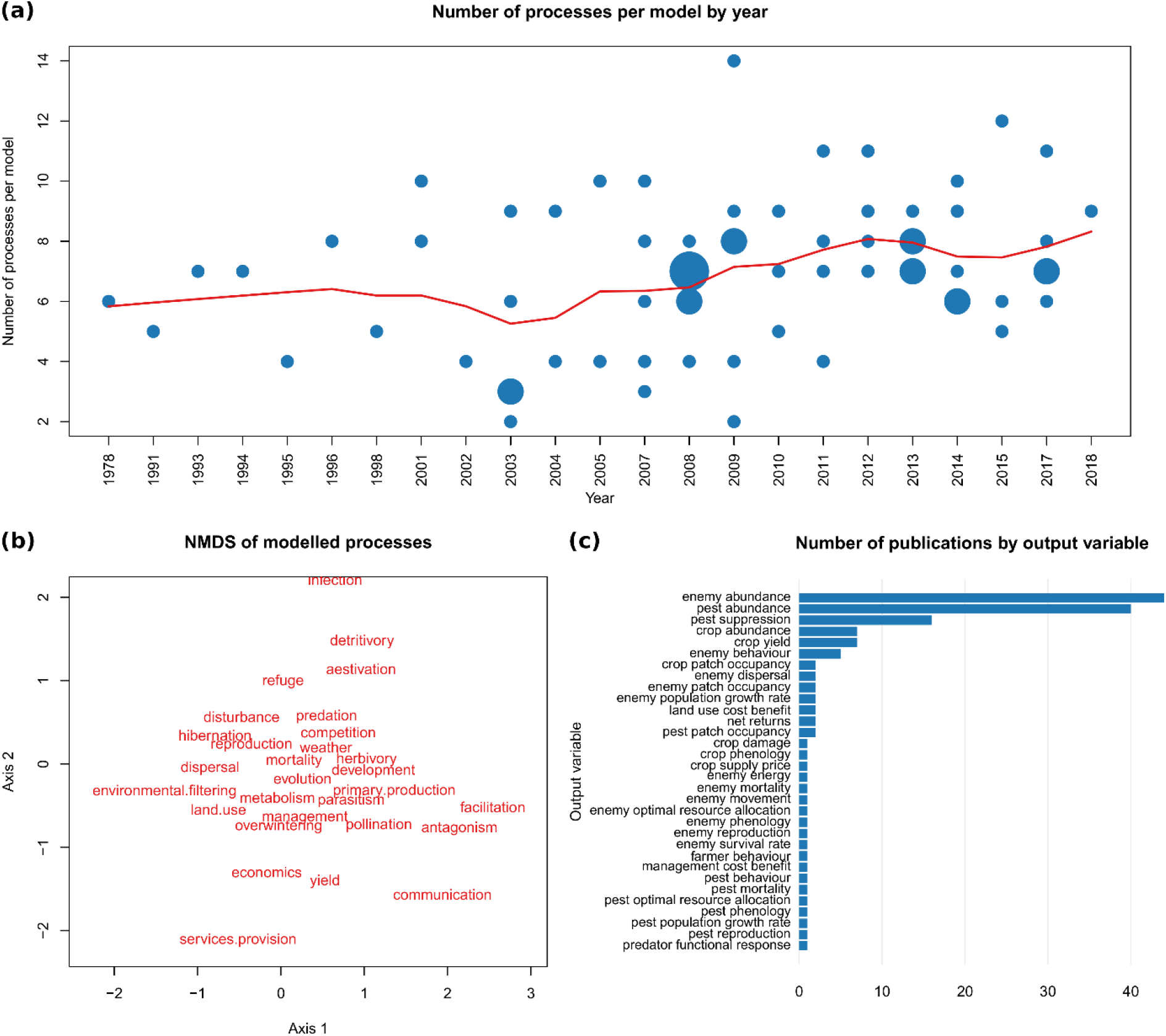
(a) Number of processes represented per reviewed mechanistic model for each publication year. Small, medium and large circles indicate 1, 2 and 3 models, respectively, with identical values. The red lowess curve was fitted to the data with a smoother span of ¼. (b) Nonmetric multidimensional scaling (NMDS) showing the associations of processes represented among reviewed mechanistic models (stress = 0.2). The closer the processes are, the more often they are represented together. (c) Number of reviewed publications having each variable as the main output of their mechanistic models. Single models may have several variables as their main output.

#### Modelling approaches

Mechanistic models of natural pest control were separated into 3 categories, representing different modelling strategies that produce models with different characteristics. Specific models are developed to represent narrowly defined crop-pest-enemy systems. Theoretical models focus on the essential features underlying the behaviour of natural pest control across systems. Conceptual models represent causal relationships that may apply to several systems without quantifying them.

Most mechanistic models represent specific crop-pest-enemy combinations (Fig. 1a), aiming to realistically represent real world systems. They typically require significant amounts of data and include the largest number of processes per model (Fig. 1b). However, they have only been developed for a few combinations of crops, pests and enemies around the world (Fig. 3a). These models originate (first author affiliation) exclusively in North America, Europe, Oceania and East Asia. The modelled systems extend to the Caribbean, South-East Asia, West and East Africa, but with a relatively small number of studies. The crop, pest and enemy organisms of these models are in many cases not specified, but there is considerable variability among the explicitly mentioned crops (Fig. 3b). Aphids are by far the most represented pests (Fig. 3c), while enemies are to a smaller extent dominated by lady beetles, wasps and spiders (Fig. 3d).

**Fig. 3.**
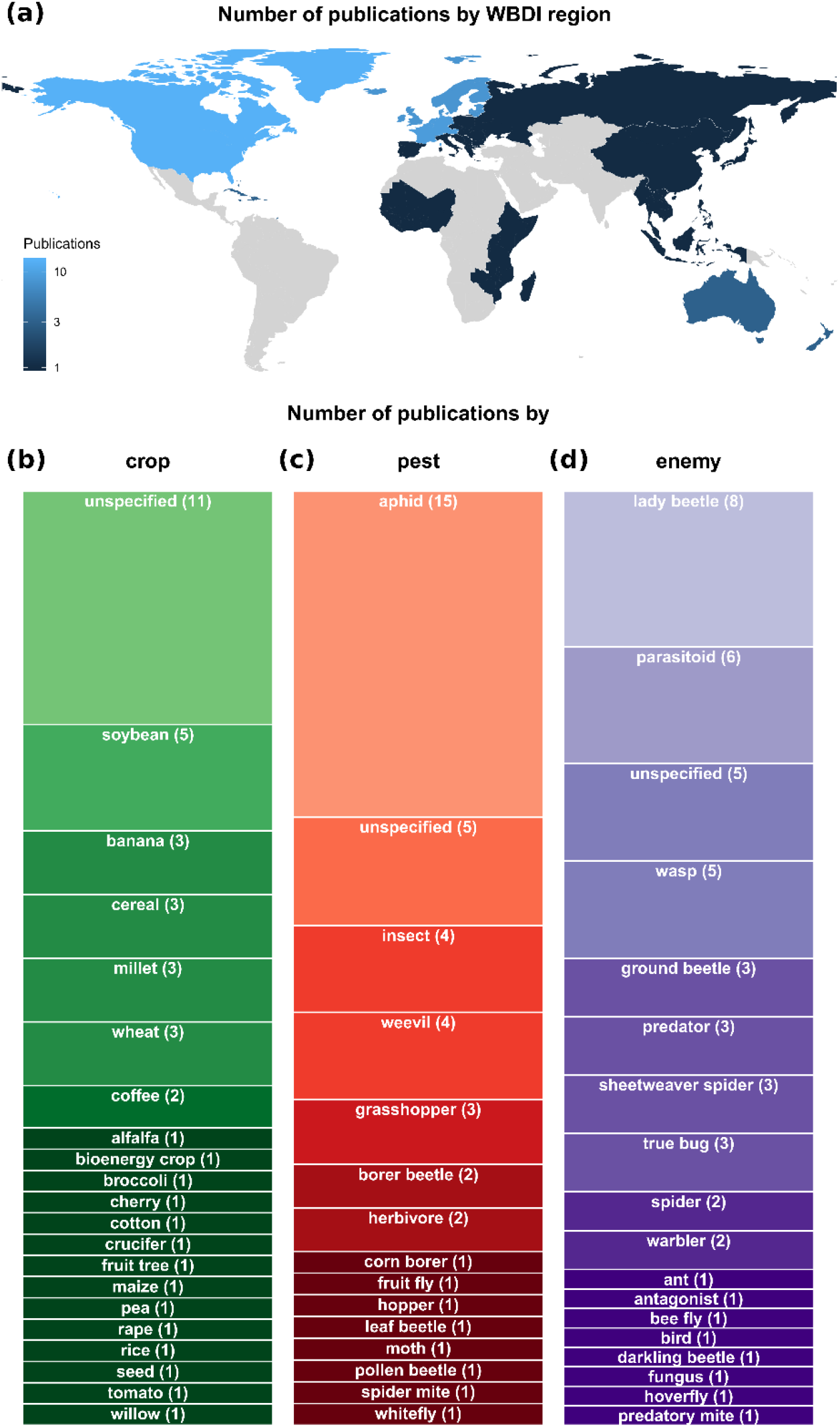
Number of reviewed publications featuring mechanistic models of specific systems (a) geographically belonging to each region defined in the World Bank Development Indicators (WBDI) and representing different (b) crop, (c) pest and (d) natural enemy organisms. Certain models did not represent all categories of organisms, others represented more than one organism in a category and, in some cases, organisms were part of the model but not identified (indicated as unspecified).

The fewer theoretical models of natural pest control (Fig. 1a) aim at producing general understanding and predictions that may apply to many systems. These models represent on average the fewest processes per model among all strategies (Fig. 1b) and they consistently approximate these processes by averaging over space. The resulting qualitative mismatch with observed phenomena, in terms of both representation and prediction, limits the ability of theoretical models to inform policy or management of natural pest control.

Conceptual models are not constrained by quantification and are able to provide a more holistic and therefore realistic system representation (Fig. 1b). However, their abstract nature typically prevents translating their representation of a system into precise responses to environmental change, hindering the comparison of model predictions with observations of a system under change. This, in turn, prevents model validation and analysis toward mechanistic understanding and reliable prediction.

#### Technical characteristics

Various techniques have been applied for mechanistic modelling of natural pest control, involving trade-offs in a model’s ability to represent different processes (or formulation flexibility), ease of analysing model output (or analytical tractability) and frugality in the use of computational resources (or algorithmic efficiency) (Table 1). Spatially explicit and stochastic representations are particularly susceptible to these trade-offs. Most models are dynamic, either continuous in time or with discrete steps. Time steps vary from 1 s to 1 yr and temporal extent varies from 1 min to 600 yr. Space is represented explicitly or implicitly by the majority of the models. Spatial resolution varies from 0.01 mm to 500 m and spatial extent from 4.5 m to 700 km.

**Table 1.**
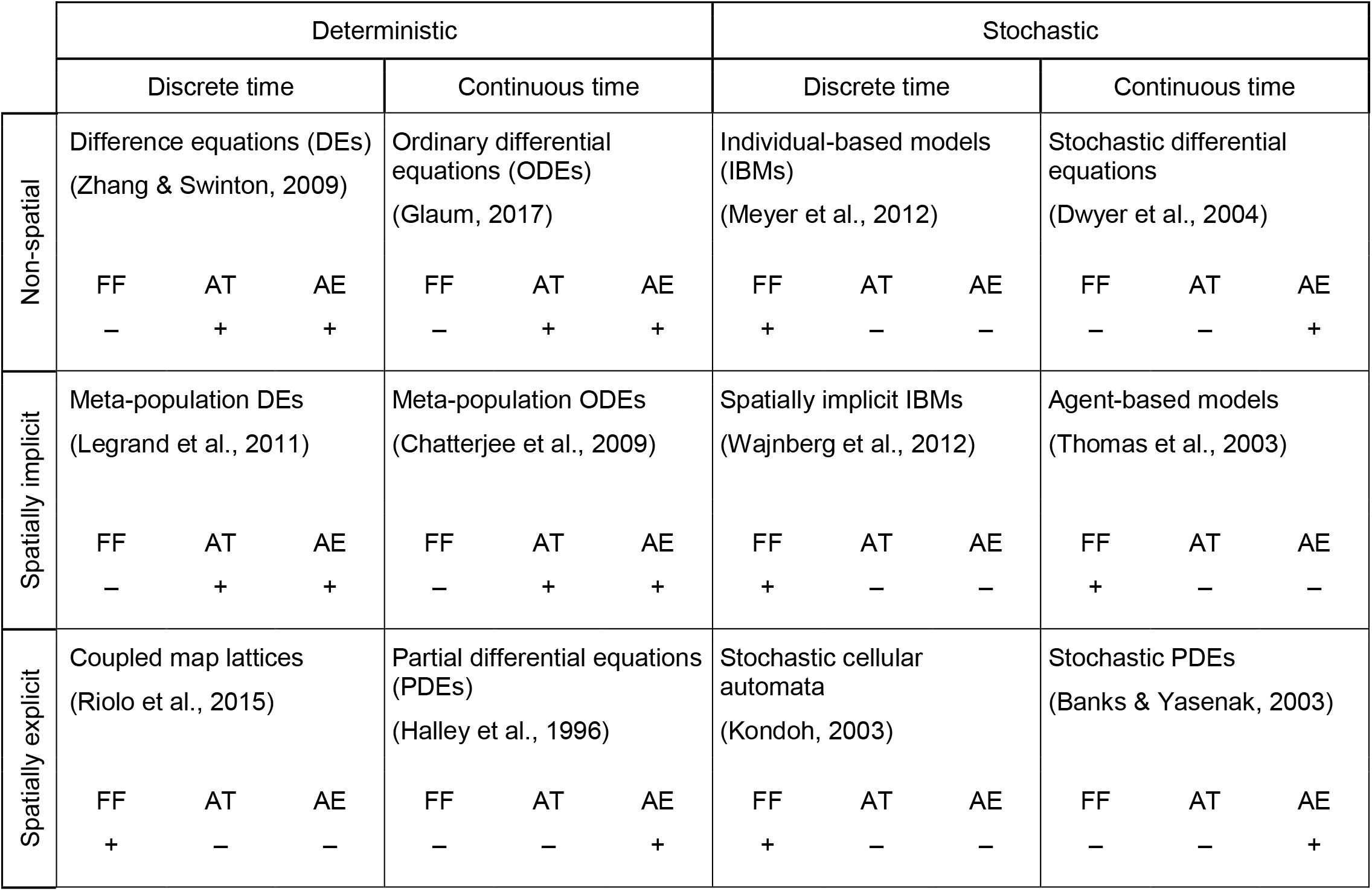
Examples of deterministic (always producing the same output given a particular input) and stochastic (including random elements) techniques used in mechanistic models of natural pest control. Technique classification is further based on the degree of explicitness in the representation of space and the distinction between discrete and continuous time representations, all viewed as key factors in modelling ecological processes. Relative strengths (+) and weaknesses (–) were assigned to each technique with respect to the following desired qualities of a modelling procedure: formulation flexibility (FF), analytical tractability (AT) and algorithmic efficiency (AE). We used the terms “individual-based” and “agent-based” to distinguish models that pool and equate individuals at each time step from models that persistently track agents independently.

The majority of mechanistic models of natural pest control represent the general ecological processes of reproduction, mortality, dispersal, environmental filtering (or species sorting) and predation along with the anthropogenic processes of land use and agricultural management. Competition, biological development, parasitism and herbivory are also represented by a considerable number of models. A few case-specific ecological as well as most socioeconomic processes form unique combinations that are represented only by a few models (Fig. 2b).

Model output has primarily comprised ecological components of natural pest control and to a smaller extent its agronomic and economic benefits, such as crop yield or net returns (Fig. 2c). The abundance of enemies, pests and, to a lesser degree, crops is the main output of the majority of mechanistic models of natural pest control. Crop yield is explicitly predicted by only 7 of the models, as more focus is put on pest suppression. A few models also predict parameters of population dynamics, as well as behavioural, physiological and economic variables related to pest control.

### Toward general predictive models

Our review of existing models of natural pest control shows a lack of realistic approaches that can provide understanding and testable predictions across agricultural landscapes. Findings agree with the hypothesis of a fundamental trade-off between generality, realism and precision among modelling strategies for biological populations (Levins, 1966). Precise and realistic models sacrifice generality by representing narrowly defined agroecosystems; general and precise theoretical models lag behind other approaches in the realism of their system representation; the few general and relatively realistic conceptual models cannot produce precise and testable predictions. Poorly studied agroecosystems in less developed regions of the world are disproportionately afflicted by this methodological gap.

The lack of general and realistic models of natural pest control can be partly addressed through qualitative mathematical modelling (Levins, 1998). The strengths of this approach, such as facilitating participatory model development and inclusion of socioeconomic factors in models of ecological systems (Fulton et al., 2015), can be of great use in predictions of natural pest control. Advances in the analysis of qualitative models (Dambacher et al., 2002, 2003) make them much more efficient than conceptual models, while retaining their strengths. Still, the lack of precision can be a significant weakness in case of non-linearities or transient dynamics, which are preferably addressed through quantitative approaches. Qualitative and quantitative models can also be developed independently for the same system. Their shared predictions are more robust against uncertainties in model formulation, a common challenge in the study of ecological systems (Levins, 1966).

The vast majority of models of natural pest control sacrifice either generality or realism and are therefore unsuited to inform management of this biologically diverse and context-sensitive phenomenon across systems. We propose that filling this methodological gap requires an approach akin to middle-range theories, providing explanations that are valid in multiple cases, but within a narrower range of conditions than grand theories (Meyfroidt et al., 2018). The number of unspecified crop-pest-enemy organisms in models of specific agroecosystems indicates some potential for generalization. This can be achieved by identifying similarities and differences that determine the responses of a variety of systems and use them to reduce the degrees of freedom that are required for multi-system representations. Studies that build consensus on drivers of behaviour across systems and identify characteristics with cross-system explanatory potential (e.g. Karp et al., 2018; Martin et al., 2019) are a key resource for this task. The result should resemble minimum realistic models, which aim to include only key variables and interactions of managed marine ecosystems (Punt & Butterworth, 1995), or the dominant processes concept, which identifies and models the most determinant hydrological processes (Grayson & Blöschl, 2001). Unlike these approaches, we argue for a framework targeted not at representing specific systems, but rather at modelling an ecological phenomenon across systems.

Existing mechanistic models of natural pest control indicate likely attributes of a general predictive framework. It appears that certain processes, including reproduction, mortality, dispersal, environmental filtering, competition and herbivory along with agricultural land use, are central to the representation of natural pest control. However, a general modelling framework should allow for context-specific flexibility with respect to the modelled processes. Even larger flexibility may be necessary in terms of temporal and spatial scales. A spatially explicit representation would be beneficial for modelling natural pest control, especially with the goal of pest management through spatial planning of agricultural landscapes. However, the representation of spatially explicit, along with stochastic, processes appears to exacerbate fundamental technical trade-offs, possibly requiring the use of novel approaches to spatial modelling. In any case, model output should expand beyond the ecological components of natural pest control, to include variables that are directly relevant to farmers, such as crop yield and economic cost-benefit measures. The resulting models will constitute context-specific generalizations that can mechanistically explain inconsistencies in the responses of natural pest control and synthesize the related socioecological knowledge. The projected increase in model development can thus be channelled toward a concerted effort to better understand and predict natural pest control around the world, further facilitating the transfer of knowledge from well-studied to poorly studied systems that share key characteristics.

